# TraPS-VarI: a python module for the identification of STAT3 modulating germline receptor variants

**DOI:** 10.1101/173047

**Authors:** Daniel Kogan, Vijay Kumar Ulaganathan

## Abstract

**Motivation:** Human individuals differ because of variations in the DNA sequences of all the 46 chromosomes. Information on genetic variations altering the membrane-proximal binding sites for signal transducer of transcription 3 (STAT3) is valuable for understanding the genetic basis of cancer prognosis and disease progression (Ulaganathan et al, 2015). In this regard, non-synonymous coding region mutations resulting in the alteration of protein sequence in the juxtamembrane region of the type I membrane proteins are biologically and clinically relevant. The knowledge of such rare cell line- and individual-specific germline receptor variants is crucial for the investigation of cell-line specific biological mechanisms and genotype-centric therapeutic approaches.

**Results:** Here we present TraPS-VarI (**Tra**nsmembrane **P**rotein **S**equence **V**ariant **I**dentifier), a python module to rapidly identify human germline receptor variants modulating STAT3 binding sites by using the genetic variation datasets in the variant call format 4.0. For the found protein variants the module also checks for the availability of associated therapeutic agents and ongoing clinical trial studies.

**Availability:** The Source code and binaries are freely available for download at https://gitlab.com/VJ-Ulaganathan/TraPS-VarI and the documentation can be found at http://traps-vari.readthedocs.io/.

**Contact:** &

**Supplementary information:** Supplementary data enclosed with the manuscript file.

## 1 Introduction

Approximately 30% of all open reading frames (ORFs) in the mammalian genome encode for membrane proteins (Krogh et al, 2001; Stevens & Arkin, 2000; Wallin & Heijne, 1998; Yildirim et al, 2007). And a large majority of all currently available therapeutic agents target membrane proteins^4^. Furthermore, it is becoming increasingly evident that variations affecting the amino acid sequence of membrane proteins might play a substantial role in disease susceptibility, disease progression and can determine therapeutic outcomes (Csaszar & Abel, 2001; Hargreaves et al, 2015; Molnar et al, 2016). Furthermore, surmounting data on genetic association of membrane protein variants with complex diseases such as autoimmune diseases, metabolic disorders, cardiovascular diseases, and cancer demands new tools for systematic approaches. Although there is no shortage of bioinformatics software catering the need of biological interpretation of genome datasets, the significance of functionally relevant variations in membrane proteins has largely remained under appreciated. Recently, we uncovered the occurrence of biologically relevant membrane-proximal STAT3 binding sites in type I membrane proteins that are capable of modulating the amplitude of STAT3 signaling within tissues and additionally altering the growth inhibition responses to certain pharmacological inhibitors (Ulaganathan et al, 2015; Ulaganathan & Ullrich, 2016). The knowledge on the frequencies of variations that either create, delete or expose such membrane-proximal STAT3 binding motifs in the general population and patient cohorts can thus significantly help clinicians to determine an effective therapeutic regimen and additionally help researchers identify and validate disease linked receptor variants. Towards this goal, we present TraPS-VarI, a python tool to rapidly identify functionally meaningful germline receptor variants by using the genome-wide genetic variation datasets.

## 2 Methods

TraPS-VarI traces genomic variants to their effects on membrane proteins using a mapping path running through nodes that include the latest human genome builds (GRCh37 and GRCh38) (Human Genome Sequencing, 2004) (MacDonald et al, 2014), Human Ensembl (Yates et al, 2016), dbSNP (Sherry et al, 2001) and Uniprot protein (2017). After determining the optimum path, the algorithm converts stepwise from genomic variant to protein substitution (**Fig. 1a**). Once the mutations on the chromosome loci are successfully converted to protein sequence alteration, the script scans the protein sequence from the end of transmembrane segment in the C-term direction until it reaches the end of 40 amino acid distances. The effect of protein substitution on the presence, absence or alteration of any membrane proximal STAT3 binding motif namely “YxxQ” (where x is any amino acid) in the 40 amino acid long membrane proximal cytoplasmic domains are reported with their position followed by the motif in parenthesis (**Fig. 1b**). Annotations are as follows, “ PRESENT”, when a motif is found but not altered by the mutation, “ABSENT”, when no motif exists, “CREATED”, when no motif is found but the mutation creates one and “DESTROYED”, when a motif is found but destroyed by the altered allele.

**Fig. 1.**
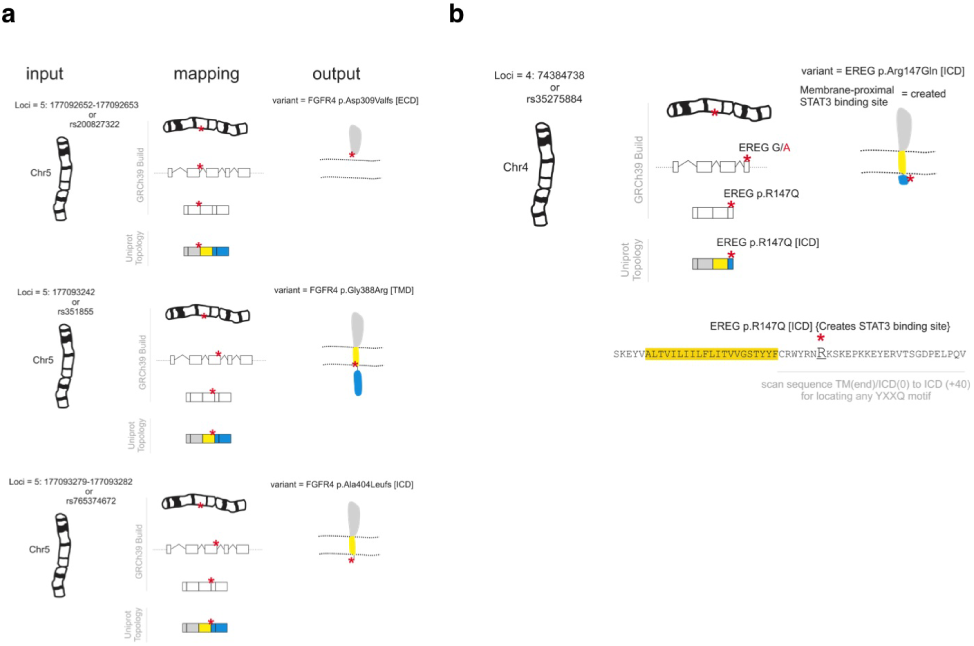
Mapping of genetic mutation altering membrane proximal STAT3 binding sites. (a) Illustration of mapping of the entries namely ‘chromosome’ and ‘position’ from the .vcf file to amino acid location in UniProt protein and (b) identification of membrane proximal STAT3 binding motif YXXQ.

## 3 Results

TraPS-VarI processes the vcf file line by line. It takes the position, matches this against coding regions in the RefSeq database. It then matches the CDS to its appropriate UniProt entry, modifies the CDS according to the mutation and retranslates the resulting CDS. The effect of the mutation is derived from the difference between those two entries. In it’s current version it only matches against UniProts main entries and not their isoforms (support for this is planned). It also looks up the position and mutation in the dbSNP dataset; if the mutation is contained there it adds the dbSNPid to the entry. It also checks the found refseq and uniprot ids against the therapeutic target database (ttd) (Zhu et al, 2012) and DrugBank database (Law et al, 2014). TraPS-Vari outputs the results in 8 columns namely, “protein id”, “protein position”, “predicted protein change”, “mutation type”, “STAT3 site”, “protein domain”, “ttd” and “drugbank”.

### 3.1 Installation

TraPS-VarI will add itself as a module to python.

#### 3.1.1 Requirements

1. Python 3.4 or newer
2. MySQLdb module for python (MySQL-python).
3. Access to a MySQL database (InnoDB engine with spatial index support - v5.7 or higher).

#### 3.1.2 Install command

Extract the package to the desired install location and run the installation script with:

> python install .py

And follow the instructions.

#### 3.1.2 Usage

python TraPSVarI.py {−p=<CHR:POS> −m=<REF/ALT>| −f=<filename> [ options ]} [-assembly=<assembly version >]

- -p=chr:pos to look up a single mutation (i.e. A/T) at the position chr:pos (only with -m)
- -m=REF/ALT mutation to look up (only in conjunction with -p)
- -f= to run the script on a file in vcf format
- -fout file to save the result to (default is input file traps vari output)
- -filter use to omit all lines from the result that do not contain a transmembrane protein mutation
- -assembly=37/38 (which assembly to use, dbSNP is only supported for 38)

The resulting file will have the protein position, protein allele change, mutation effect and the motif changes, hits in the ttd database and the DrugBank database appended as columns. Additionally, snippets that are readily integrable in the workflow of TraPS-VarI module are available under https://github.com/VJ-Ulaganathan.

## Acknowledgements

The authors thank Prof. Dr. Axel Ullrich for supporting this work and Viet Nuyen for technical assistance.

## Conflict of Interest

The authors declare no conflict of interests.

---

**Supplementary_Figure. 1.**
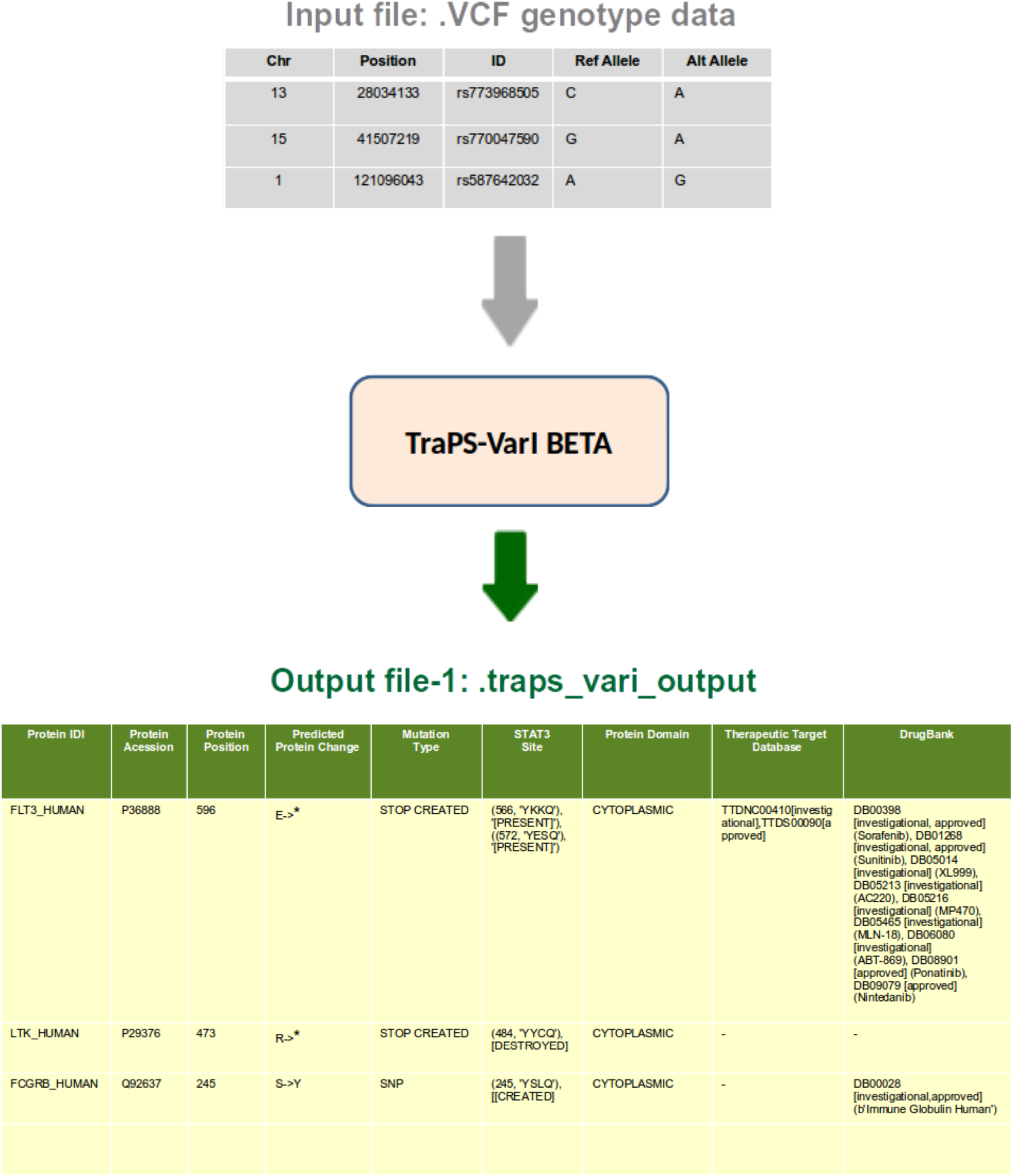
Schematic diagram describing the workflow of TraPS-VarI python module

